# Pathogenic variants associated with speech/cognitive delay and seizures affect genes with expression biases in excitatory neurons and microglia in developing human cortex

**DOI:** 10.1101/2024.07.01.601597

**Authors:** Jeffrey B. Russ, Alexa C. Stone, Kayli Maney, Lauren Morris, Caroline F. Wright, Jillian H. Hurst, Jennifer L. Cohen

## Abstract

**Background & Objective:** Congenital brain malformations and neurodevelopmental disorders (NDDs) are common pediatric neurological disorders and result in chronic disability. With the expansion of genetic testing, new etiologies for NDDs are continually uncovered, with as many as one third attributable to single-gene pathogenic variants. While our ability to identify pathogenic variants has continually improved, we have little understanding of the underlying cellular pathophysiology in the nervous system that results from these variants. We therefore integrated phenotypic information from subjects with monogenic diagnoses with two large, single-nucleus RNA-sequencing (snRNAseq) datasets from human cortex across developmental stages in order to investigate cell-specific biases in gene expression associated with distinct neurodevelopmental phenotypes.

**Methods:** Phenotypic data was gathered from 1) a single-institution cohort of 84 neonates with pathogenic single-gene variants referred to Duke Pediatric Genetics, and 2) a cohort of 4,238 patients with neurodevelopmental disorders and pathogenic single-gene variants enrolled in the Deciphering Developmental Disorders (DDD) study. Pathogenic variants were grouped into genesets by neurodevelopmental phenotype and geneset expression across cortical cell subtypes was compared within snRNAseq datasets from 86 human cortex samples spanning the 2nd trimester of gestation to adulthood.

**Results:** We find that pathogenic variants associated with speech/cognitive delay or seizures involve genes that are more highly expressed in cortical excitatory neurons than variants in genes not associated with these phenotypes (Speech/cognitive: p=2.25×10^-7^; Seizures: p=7.97×10^-12^). A separate set of primarily rare variants associated with speech/cognitive delay or seizures, distinct from those with excitatory neuron expression biases, demonstrated expression biases in microglia. We also found that variants associated with speech/cognitive delay and an excitatory neuron expression bias could be further parsed by the presence or absence of comorbid seizures. Variants associated with speech/cognitive delay *without* seizures tended to involve calcium regulatory pathways and showed greater expression in extratelencephalic neurons, while those associated with speech/cognitive delay *with* seizures tended to involve synaptic regulatory machinery and an intratelencephalic neuron expression bias (ANOVA by geneset p<2×10^-16^).

**Conclusions:** By combining extensive phenotype datasets from subjects with neurodevelopmental disorders with massive human cortical snRNAseq datasets across developmental stages, we identified cell-specific expression biases for genes in which pathogenic variants are associated with speech/cognitive delay and seizures. The involvement of genes with enriched expression in excitatory neurons or microglia highlights the unique role both cell types play in proper sculpting of the developing brain. Moreover, this information begins to shed light on distinct cortical cell types that are more likely to be impacted by pathogenic variants and that may mediate the symptomatology of resulting neurodevelopmental disorders.

## INTRODUCTION

Neurodevelopmental disorders (NDDs) broadly encompass both macroanatomic congenital brain malformations, as well as microcircuit and molecular disorders of development, such as autism spectrum disorder, epilepsy, and developmental delay. Although any individual disorder may be somewhat rare, their cumulative prevalence is high and often results in significant, chronic disability for many pediatric patients. Depending on precise definitions and screening techniques, congenital brain malformations are detected in approximately 0.2 to 10 per 1000 pregnancies,^1–5^ while anywhere between 3 and 170 per 1000 children have a NDD.^6–10^ Co-occurrence of multiple neurodevelopmental symptoms is common in NDDs, since primary cortical malformations, hydrocephalus, or perinatal vascular injury can result in secondary seizures, cerebral palsy, or developmental delay.^6,11–13^ In fact, high rates of neurodevelopmental comorbidities have made it challenging to pinpoint the distinct genetic and cellular pathophysiology of individual neurodevelopmental symptoms and untangle them from the comorbid pathophysiology of co-existing neurodevelopmental condtions.^14^

With increasingly available comprehensive genetic testing, our ability to uncover genetic etiologies for many patients with NDDs is continually improving.^14–19^ Large, national studies, such as the Deciphering Developmental Disorders (DDD) study, have begun depositing highly detailed, standardized phenotypic information into accessible databases and linking them with genome-wide data and specific genetic diagnoses.^16, 20^ Despite an enhanced understanding of genotype-phenotype relationships for many genetic forms of NDDs, our understanding of the nuanced cortical pathophysiology that likely mediates the symptoms of these conditions remains sparse. Novel strategies are required to begin untangling the interplay between pathogenic genetic variants, their effect on highly specific cortical cell types, and the subsequent cell-specific dysfunction that underlies NDDs.

In parallel with the rise of enhanced clinical genetic testing, there has also been rapid advancement in our ability to probe gene expression at a single-cell resolution, yielding ever expanding single-cell transcriptomic atlases of human cortical development.^21–24^ Through this work, a hierarchy of distinct cortical cell types defined by their transcriptomic signatures has emerged, allowing us to probe the expression levels of tens of thousands of genes in individual cells from human postmortem cortex at increasing scale.^25, 26^ Previous studies have used this information to evaluate the convergence of autism-associated genes^22, 24, 27^ or congenital hydrocephalus-related genes^28^ in human cortical transcriptomic data. However, these studies evaluate only a single neurodevelopmental phenotype, include fewer than two hundred disease-related genes, and map them within transcriptomic datasets of fewer than 20,000 cells. Given the potentially significant overlap between genes underlying multiple NDDs and the availability of large, single-nucleus and single-cell gene expression databases, there is an opportunity to uncover patterns of gene expression within specific cortical cell types that contribute to neurodevelopmental symptoms across disorders.

To investigate whether genes associated with certain neurodevelopmental phenotypes exhibit cortical cell-specific expression biases, we combined the rich genotype-phenotype data from two large cohorts of pediatric subjects with NDDs with the wealth of gene expression data from two massive single-nucleus RNA-sequencing (snRNAseq) datasets from postmortem human cortex samples collected across developmental stages. Combining these datasets, we uncover expression biases in both excitatory cortical neurons and microglia for genes associated with speech/cognitive delay and/or seizures.

## MATERIALS AND METHODS

### Collection and curation of Duke Pediatric Genetics data

Retrospective chart review was performed on a cohort of 334 neonates referred from the Duke Intensive Care Nursery to Pediatric Genetics for any reason, and who subsequently had at least one pathogenic variant or variant of uncertain significance (VUS) on molecular genetic testing. Data collected included subject sex; age at last follow up by Pediatric Neurology, Special Infant Care Clinic, or Developmental Pediatrics; and the presence or absence of neurodevelopmental symptoms or NDDs. Ages at last follow up date ranged from five days old to 7.75 years old. The binary presence or absence of symptoms was coded as positive if symptomatology was documented at any age before their most recent follow up in the above clinics. Autism spectrum disorder, seizures, motor delay, and speech or cognitive delay were coded as a binary variable - present or absent. Seizures were coded as present if patients had a history of any type of seizure and was not restricted to a diagnosis of epilepsy. Speech or cognitive delay were combined into one variable since more detailed information necessary to separate these two symptoms was not consistently available. All severities of motor delay or speech or cognitive delay were counted as those conditions being present. Structural brain malformations were first coded as present or absent, with relevant details included in a free-text form. An experienced Pediatric Neurologist (J.B.R.) reviewed each positive case and coded free text answers as binary variables (present or absent) for midline disorders, malformations of cortical development, corpus callosum dysgenesis, hydrocephalus/ventriculomegaly, or vascular disorders. Ventriculomegaly due to hydrocephalus was not considered separately from ventriculomegaly from other causes since information on the etiology was not standardized or consistently available for all subjects. Vascular disorders included any history of ischemic brain injury, intracranial hemorrhage, or vascular malformation and were combined into one binary variable.

Information on genetic diagnoses was collected from any available clinical genetic testing, including chromosomal microarray, targeted molecular next-generation sequencing (NGS) or NGS gene panels, exome sequencing, or genome sequencing. Variant information was documented and the predicted pathogenicity, as determined and reported by a clinical diagnostic laboratory, was documented as pathogenic (including likely pathogenic), uncertain significance, or benign (including likely benign). Variants for downstream analysis in the snRNAseq data were restricted to single nucleotide changes or small deletions/duplications that affected single genes and were predicted to be pathogenic (or likely pathogenic). Overall, the dataset included 92 pathogenic variants from 84 subjects in 60 unique genes.

Study procedures were reviewed and declared exempt by the Duke University Institutional Review Board (Pro00106469).

### Collection and curation of DDD study data

The DDD study protocol has been extensively described elsewhere.^20^ We obtained information on pathogenic variants detected in probands with NDDs from the Deciphering Developmental Disorders (DDD) study, as reported in Wright et al., 2023,^16^ from the European Genome-Phenome Archive (https://ega-archive.org/studies/EGAS00001000775). Genotype information (EGAD00001010137) was linked with individual-level phenotype information (EGAD00001004388). Analyses were restricted to variants with single nucleotide changes or small deletions/duplications that affected single genes and were predicted to be pathogenic or likely pathogenic, as classified by referring clinical teams in the DDD study. Overall, the dataset included 4,621 pathogenic variants from 4,238 subjects in 862 unique genes. To code phenotype information based on the binary presence or absence of specific neurodevelopmental symptoms, Human Phenotype Ontology (HPO) terms were grouped as outlined in Supplemental Table 1. If one or more phenotype associated HPO terms was present, the neurodevelopmental phenotype was coded as present.

### Human cortex snRNAseq datasets

Raw snRNAseq data from Velmeshev et al, 2023^26^ was acquired from the UCSC Cell Browser repository (https://cells.ucsc.edu/?bp=brain&org=Human+(H.+sapiens)&ds=pre-postnatal-cortex), using the raw count matrix from “All Cells.” Detailed cell type metadata was obtained from the meta.tsv files available under “Excitatory Neurons,” “Glial Cells,” “Interneurons,” “Microglial Cells,” and “Vascular Cells.” Metadata cell type information was merged with the metadata from “All Cells” by cell barcode.

Raw snRNAseq data from Herring et al., 2022^25^ was acquired from the Lister lab Google bucket (http://brain.listerlab.org). The raw count matrix “RNA-all_full-counts-and-downsampled-CPM.h5ad” was obtained and converted to a Seurat object using the anndata package (version 0.7.5.6) in R.

Both raw snRNAseq datasets were then reconstructed using the Seurat package (version 5.0.0) in R. Normalization, variable feature selection, and scaling were all performed via the SCTransform function. Principal component analysis (PCA) was performed using RunPCA and UMAP coordinates were obtained by performing RunUMAP on the first 30 dimensions. UMAP clustering was performed using FindNeighbors(dims=1:30) and FindClusters(resolution=0.2) to ensure that cluster boundaries were similar to the originally published objects; however, the original metadata was retained for cell type annotation, which showed faithful reclustering by cell type.

### Measuring variant gene expression biases in human cortical snRNAseq datasets

Given the wide variability in magnitude of endogenous gene expression across cell type-specific or developmental stage-specific populations for NDD-associated genes included in this study, a standardized metric of relative gene enrichment was required. Thus, FindAllMarkers was performed on the “lineage” and “age_range” idents from the Velmeshev dataset or on the “cell_class” and “stage_id” idents from the Herring dataset using the following parameters: assay=“RNA,” only.pos=TRUE, min.pct=0.25, logfc.threshold=0.25. For any given gene that was significantly enriched (adjusted pval<0.05) in a cell type- or developmental stage-specific subpopulation, this provided a log2-fold change in expression level above the cumulative expression level within all other subpopulations. It also yielded a calculation of the percent of cells in a subpopulation of interest that expressed the gene as compared to the cumulative percentage of cells from all other subpopulations that express the gene. Each gene associated with a pathogenic variant in each phenotype geneset was then amended with a relative expression level and a relative percent expression for each cell type (i.e., Excitatory) or developmental stage (i.e., 2^nd^ Trimester). Since metrics are only calculated for subpopulation-enriched genes, the relative expression level and percent expression were imputed as zero for non-enriched genes.

Gene expression within each cell type was categorized as highly expressed or not highly expressed based on whether that gene was expressed in greater than 25% of cells of that cell type (pct1>0 after thresholding min.pct=0.25 during the FindAllMarkers step). We then used this variable to calculate the percent of subjects positive for a specific phenotype based on whether their diagnostic gene was or was not highly expressed in each cell type.

To parse excitatory neuron biased genes associated with speech/cognitive delay and/or seizures, we first classified genes based on whether they were associated with seizures in any subject and whether they were associated with speech/cognitive delay in any subject. Genes in which pathogenic variants were not associated with speech/cognitive delay in any subject were categorized as “No Speech/Cognitive Delay.” Genes that were associated with speech/cognitive delay but had a log2-fold change in excitatory neurons less than 0.25 and/or was expressed in less than 25% of excitatory neurons were categorized as “Non-Excitatory Speech/Cognitive Delay.” Genes that showed a log2-fold enrichment greater than 0.25 in excitatory neurons and/or were expressed in greater than 25% of excitatory neurons, were associated with speech /cognitive delay, but were *not* associated with seizures in any subject were classified as “Excitatory Speech/Cognitive Delay without Seizures.” Finally, genes that showed a log2-fold enrichment greater than 0.25 in excitatory neurons and/or were expressed in greater than 25% of excitatory neurons and were also associated with both speech/cognitive delay and seizures were classified as “Excitatory Speech/Cognitive Delay with Seizures.” Mean log2-fold change and percent expression of variant genes were then plotted across these four categories. Log2-fold change was used to compare relative expression across the four new geneset categories by excitatory neuron subtype, as previously annotated,^26^ or by developmental stage at which the post-mortem sample was collected.

Gene Ontology (GO) term enrichment was performed using the enrichGO function in the clusterProfiler package (version 4.10.0) in R, as mapped to human gene expression data (OrgDb = org.Hs.eg.db), and the top ten results with the lowest p-values were plotted. An aggregate geneset expression score for each of the four excitatory speech/cognitive delay and/or seizure categories were calculated for each individual cortical cell using the AddModuleScore_UCell function from the UCell package (version 2.6.1) in R.^29^ The UCell score for each cell was then plotted against its excitatory lineage pseudotime score, a proxy for its developmental stage, as previously published.^26^

To calculate enrichment of a given phenotype-associated geneset in the “gene trend” categories defined in Herring et al., 2022, we performed hypergeometric testing across gene trend annotations as defined in Herring et al. 2022 Table S3 (Supplemental Table 2).^25^

Genes associated with speech/cognitive delay and/or seizures that were enriched in microglia were subdivided into narrow or broad microglial enrichment based on whether they were expressed in less than or greater than five percent of non-microglial cells, respectively.

### Programming, Graphs, and Statistical Analysis

All analyses were performed using R/RStudio (version 2023.09.1+494). snRNAseq data was analyzed using Seurat (version 5.0.0).^30^ Feature plots were generated using the FeaturePlot command in Seurat. Graphs were generated using ggplot2 (version 3.4.4) and gplots (version 3.1.3) in R. Venn diagrams were generated with the VennDiagram (version 1.7.3) package in R. A Wilcoxon rank sum test was used to compare log2-fold change and percent expression across two conditions (i.e., the presence or absence of a neurodevelopmental phenotype). Chi-squared testing was used to compare categorical variables (i.e., the presence or absence of seizures versus the presence or absence of expression in excitatory neurons). ANOVA was used to compare continuous dependent variables across categorical independent variables followed by Tukey testing of pairwise comparisons corrected for multiple comparisons. Hypergeometric testing was used to calculate the probability of geneset enrichment within “gene trend” categories defined by Herring et al, 2022.^25^ Bonferroni correction for multiple comparisons was utilized for all Wilcoxon and hypergeometric tests. Significance was set at an adjusted p-value <0.05.

## RESULTS

### Subject datasets

For this study, we utilized two different pediatric neurodevelopmental subject datasets summarized in Table 1: 1) a single-institution dataset from Duke University of neonates referred to Pediatric Genetics from the intensive care nursery for any genetic concern whose molecular genetic testing yielded at least one pathogenic variant or VUS, and 2) a dataset from the Deciphering Developmental Disorders (DDD) study,^16^ a national United Kingdom/Ireland study of probands with severe NDDs. The advantage of the single-institution dataset is that it contains data from all-comers with data collection beginning in the neonatal period, regardless of their eventual neurodevelopmental status. This design provides a less selective dataset that begins in the neonatal period and captures a higher proportion of subjects with congenital brain malformations. The advantage of the DDD dataset is the standardized phenotype characterization and a cohort size that dramatically increases our power to detect cell type expression biases by phenotype.

**Table 1:**
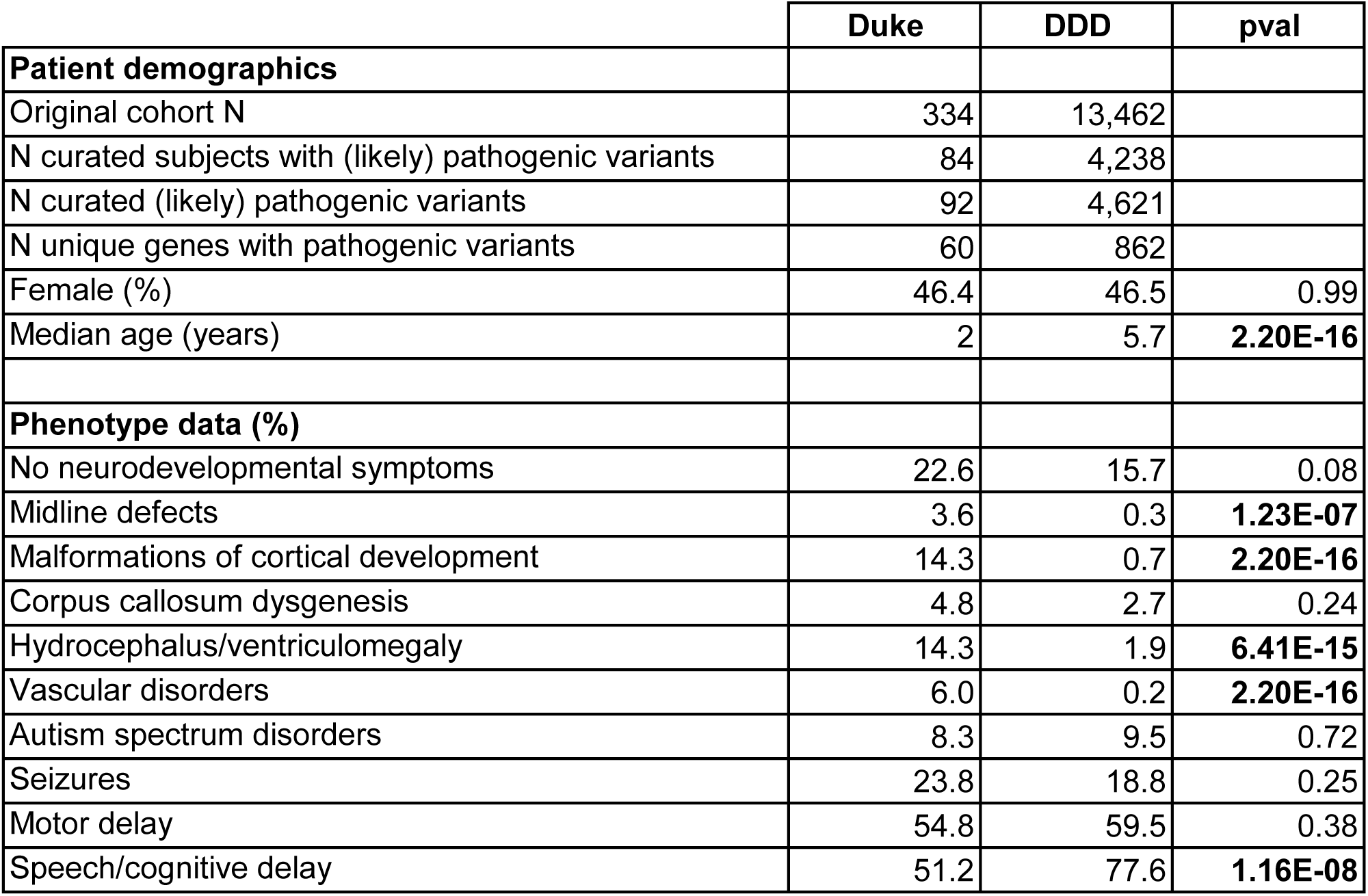
Duke versus DDD subject demographics and NDD prevalence.

The Duke Pediatric Genetics dataset initially contained 334 subjects with 488 aneuploidies, copy number variations (CNVs), or monogenic variants identified through genetic testing, at least one of which was pathogenic or of uncertain significance for each subject. We distilled this initial cohort to 84 subjects with 92 pathogenic single gene variants. Just under half of this curated cohort was female (n=39, 46.4%) and the median age at the time of phenotypic data collection was two years. Keeping in mind that individual subjects can be included in more than one phenotypic category, 3.6% of subjects had midline cerebral defects, 14.3% had malformations of cortical development (MCDs), 4.8% had corpus callosum dysgenesis (CCD), 14.3% had hydrocephalus or ventriculomegaly, 6% had vascular disorders, 8.3% had autism spectrum disorder (ASD), 23.8% had seizures, 54.8% had motor delay of any severity, and 51.2% had speech or cognitive delay of any severity. There were also 19 subjects (22.6%) who had pathogenic single gene variants in the absence of neurodevelopmental symptoms.

The DDD dataset contained 13,462 subjects with neurodevelopmental phenotypes. Of these, 4,238 subjects (31.5%) had 4,621 pathogenic or likely pathogenic single-gene variants classified by the probands’ referring clinical teams. Of this curated cohort, 1,970 were female (46.5%) and the median age at the time of recruitment and phenotypic data collection was 5.7 years old. 0.3% of the DDD cohort had midline cerebral defects, 0.7% had MCD, 2.7% had CCD, 1.9% had hydrocephalus or ventriculomegaly, 0.2% had vascular disorders, 9.5% had ASD, 18.8% had seizures, 59.5% had motor delay of any severity, and 77.6% had speech or cognitive delay of any severity. The DDD dataset also included 664 subjects (15.7%) who had pathogenic single gene variants without neurodevelopmental symptoms but with other non-neurological developmental phenotypes.

The phenotypic composition of the Duke Pediatric Genetics and DDD datasets did not differ by sex (p=0.99), or in the prevalence of CCD (p=0.24), ASD (p=0.72), seizures (p=0.25), motor delay (p=0.38), or non-NDD cases (p=0.08). The median age at the time of last follow up was significantly younger in the Duke dataset (p=2.20×10^-16^), and there was a significantly higher prevalence of midline cerebral defects (p=1.23×10 ^-7^), MCD (p=2.20×10^-16^), hydrocephalus/ventriculomegaly (p=6.41×10^-16^), and vascular disorders (p=2.20×10^-16^) compared to the DDD dataset, likely reflecting the enrollment of subjects from the neonatal stage in the Duke cohort. The prevalence of speech/cognitive delay was higher in the DDD cohort (p=1.16×10^-8^).

The curated Duke and DDD cohorts were then combined, and pathogenic variants associated with each phenotype (Supplemental Table 3) were grouped into “genesets” (Supplemental Table 4) for testing of cortical cell type expression biases by phenotype (Figure 1A). To maintain an unbiased approach, capture phenotypic variation and complexity, and to ensure the inclusion of rare phenotypes, we grouped all genes associated with a given phenotype into the geneset for that phenotype, regardless of whether direct causation has been previously established for a given genotype-phenotype combination. Similarly, to avoid over-curation and to maintain a dataset representative of the full spectrum of phenotypes across a large population, in some circumstances, a pathogenic variant in a gene known to cause a neurodevelopmental phenotype was still included in our “negative” geneset if a subject with that variant had no neurodevelopmental symptoms. As a result, some genes were included across multiple genesets (including some genes that ended up in both the negative geneset and a positive phenotypic geneset if there were mixed phenotypes in the cohort).

**Figure 1:**
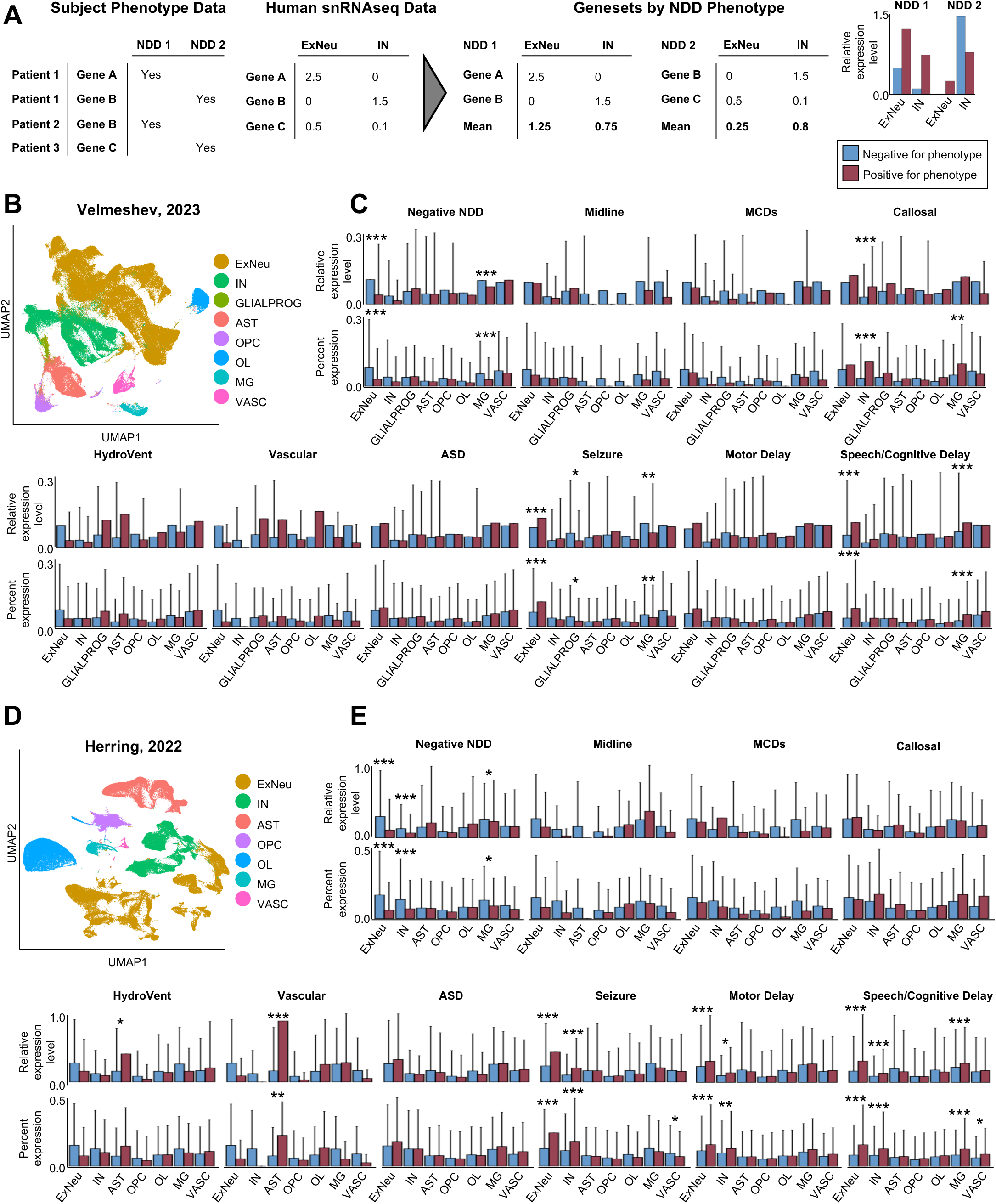
Pathogenic variants associated with NDDs affect genes with cell-specific expression biases in human cortex. **(A)** Schematic example of the data analysis pipeline used in this study. Patients with pathogenic variants in genes A, B, or C were coded on the binary presence of absence of NDD phenotype (1 or 2) and grouped into NDD-associated genesets. Separately, transcriptomic data from human cortex was analyzed for relative enrichment of each gene in each cortical cell subtype. These datasets were then merged to provide a relative expression metric for each gene within each cortical cell type and NDD geneset, which is then plotted for all genes associated with the phenotype (red) versus all genes not associated with the phenotype (blue). **(B)** UMAP of 348,049 nuclei from Velmeshev et al., 2023 with originally assigned cell type annotations. **(C)** Mean relative expression level (top; log2 fold change in expression compared to all other cell types) and mean percent expression (bottom; percent of cell type expressing genes of interest) demonstrate cell type-specific expression biases in human cortex. **(D)** UMAP of 153,473 nuclei from Herring, et al., 2022 with originally assigned cell type annotations. **(E)** Mean relative expression level (top) and mean percent expression (bottom) reproduce many of the cell type-specific expression biases observed in **(C)**. In particular, genes associated with seizures and speech/cognitive delay demonstrate higher relative expression and higher percent expression in excitatory neurons in both datasets, and genes associated with speech/cognitive delay also demonstrate an expression bias in microglia in both datasets **(C,E)**. Red columns = genes associated with the phenotype; blue columns = genes not associated with the phenotype. Abbreviations: NDD = neurodevelopmental disorder; Midline = midline defects; MCDs = malformations of cortical development; Callosal = callosal abnormalities; HydroVent = hydrocephalus/ventriculomegaly; Vascular = vascular disorders, hemorrhage, or stroke; ASD = autism spectrum disorder; ExNeu = excitatory neurons; IN = inhibitory neurons; GLIALPROG = glial progenitors; AST = astrocytes; OPC = oligodendrocyte precursor cells; OL = oligodendrocytes; MG = microglia; VASC = endothelial cells/pericytes; UMAP = uniform manifold approximation and projection. * = p<0.05; ** = p<0.01; *** = p<0.001.

### Genes with pathogenic variants associated with several neurodevelopmental phenotypes exhibit expression biases in distinct cortical cell types

To investigate whether pathogenic variants associated with specific neurodevelopmental phenotypes involved genes that exhibit cortical cell type expression biases, we utilized two recently published, publicly available snRNAseq datasets from human cortex across development.^25,26^ The Velmeshev, 2023 transcriptomic dataset includes 348,049 nuclei from human cortical specimens from 60 individuals ranging in developmental stage from second trimester gestation to adulthood (Figure 1B).^26^ To ensure reproducibility, we validated our findings in 153,473 nuclei from 26 subjects spanning the same developmental stages, collated from Herring, 2022 (Figure 1D).^25^ Maintaining the original cell annotations from both studies, we then analyzed 1) the relative expression level (defined as the fold change of gene expression in a given cell type as compared to its cumulative expression in all other cell types), and 2) the breadth of expression within each cortical cell class (defined as the percent of cells from each cell type expressing a given gene) for each phenotypic geneset (Figure 1C,E).

After correcting for multiple comparisons, we found several expression biases of phenotype-related genesets within cortical cell classes that were replicated across both human cortical snRNAseq datasets (Figure 1C,E). Most prominently, genes associated with speech/cognitive delay and genes associated with seizures demonstrated higher relative expression in excitatory cortical neurons and were also expressed in a larger percentage of cortical excitatory neurons than genes not associated with these phenotypes. Genes associated with speech/cognitive delay also showed higher relative expression and were expressed in a higher percentage of microglia than those not associated with speech/cognitive delay. In contrast, microglial expression was significantly lower for genes associated with seizures in the Velmeshev, 2023 dataset, though this was not reproduced in the Herring, 2022 dataset.

Sub-analyses demonstrated that although relative patterns of gene expression were similar between the Duke and DDD datasets, significance was largely driven by the DDD dataset (Supplemental Figure 1), which is not surprising given the much larger cohort size. Additionally, microglial enrichment of genes associated with speech/cognitive delay appeared driven predominantly by rare variants, as this effect was lost after excluding variants observed in fewer than five subjects (Supplemental Figure 2). Excitatory neuron enrichment was not affected by the exclusion of rare variants.

We focused the remainder of our analysis on the robust effects that were replicable in both snRNAseq datasets, which included excitatory enrichment in genes associated with speech/cognitive delay and seizures. We also focus on microglial enrichment of genes associated with speech/cognitive delay, despite the larger impact of rare variants on this finding. Given the larger number of nuclei and the larger patient sample, subsequent analyses were performed using the Velmeshev, 2023 dataset.^26^

We next examined the frequency of a phenotype based on whether associated genes were expressed in greater than 25% of each cell type (“high cell type expression”). This converse analysis yielded very similar results (Figure 2). Subjects whose pathogenic variants affected genes with high excitatory neuron expression were significantly more likely to have speech/cognitive delay and/or seizures than subjects whose variants affected genes that are not highly expressed in excitatory neurons. The opposite was true for negative NDD subjects whose variants were more likely to have low excitatory neuron expression. Subjects whose variants were enriched in microglia had significantly higher rates of speech/cognitive delay but lower rates of seizures. Subjects with a negative NDD phenotype were significantly likely to have variants in genes with low microglial expression. Finally, subjects with variants in genes highly expressed in interneurons or microglia had significantly higher rates of CCD, and those whose variants affected genes expressed in glial progenitors (GLIALPROG) had a small but significant decreased likelihood of having seizures.

**Figure 2:**
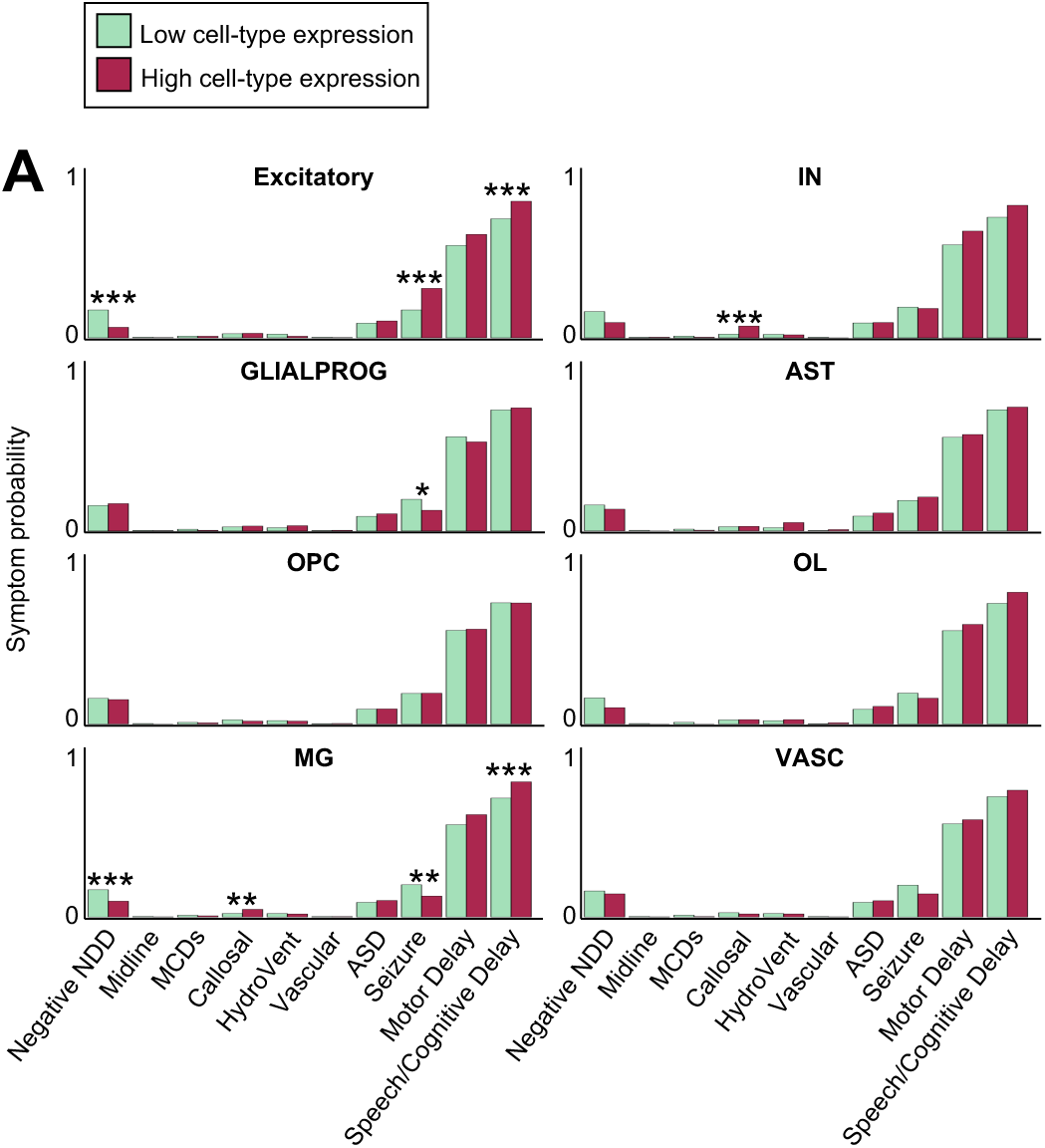
Likelihood of neurodevelopmental phenotypes by cell-specific gene expression. Genes associated with pathogenic variants were binned as “Low cell-type expression” (green) or “High cell-type expression” (red) based on whether they were expressed in fewer or more than 25% of cells of a given cell type, respectively. The probability of subjects presenting with each NDD phenotype was then calculated for genes that have low or high expression in each cortical cell type. Subjects with pathogenic variants in genes that have high excitatory neuron expression are significantly less likely to have a negative phenotype and significantly more likely to have seizures or speech/cognitive delay. Subjects with variants in genes that have high microglial expression are more likely to have speech/cognitive delay, but less likely to have seizures or a negative phenotype. Abbreviations: NDD = neurodevelopmental disorder; Midline = midline defects; MCDs = malformations of cortical development; Callosal = callosal abnormalities; HydroVent = hydrocephalus/ventriculomegaly; Vascular = vascular disorders, hemorrhage, or stroke; ASD = autism spectrum disorder; ExNeu = excitatory neurons; IN = inhibitory neurons; GLIALPROG = glial progenitors; AST = astrocytes; OPC = oligodendrocyte precursor cells; OL = oligodendrocytes; MG = microglia; VASC = endothelial cells/pericytes. * = p<0.05; ** = p<0.01; *** = p<0.001.

### Subjects with pathogenic variants in excitatory neuron-enriched genes demonstrate two distinct neurodevelopmental phenotypes

Given the highly significant and reproducible expression bias of genes associated with speech/cognitive delay and seizures in excitatory cortical neurons in both snRNAseq datasets (Figure 1), we first investigated the relationship between these two phenotypic genesets and found that genes associated with seizures were a subset of genes associated with speech/cognitive delay (Figure 3A). Stated differently, we observed two subsets of genes: 1) those associated with speech/cognitive delay *and* seizures, versus 2) those associated only with speech/cognitive delay *without* seizures. 43% of all genes associated with speech/cognitive delay were also associated with seizures, while the remaining 57% were not associated with seizures. Only 20 genes (6%) associated with seizures from the full dataset were not associated with speech/cognitive delay (not shown). Restricting this analysis only to genes that are widely expressed in excitatory cortical neurons, 65% of excitatory genes associated with speech/cognitive delay were also associated with seizures, while the remaining 35% were not associated with seizures (Figure 3A). No excitatory-enriched genes were associated with seizures without speech/cognitive delay (Figure 3A).

**Figure 3:**
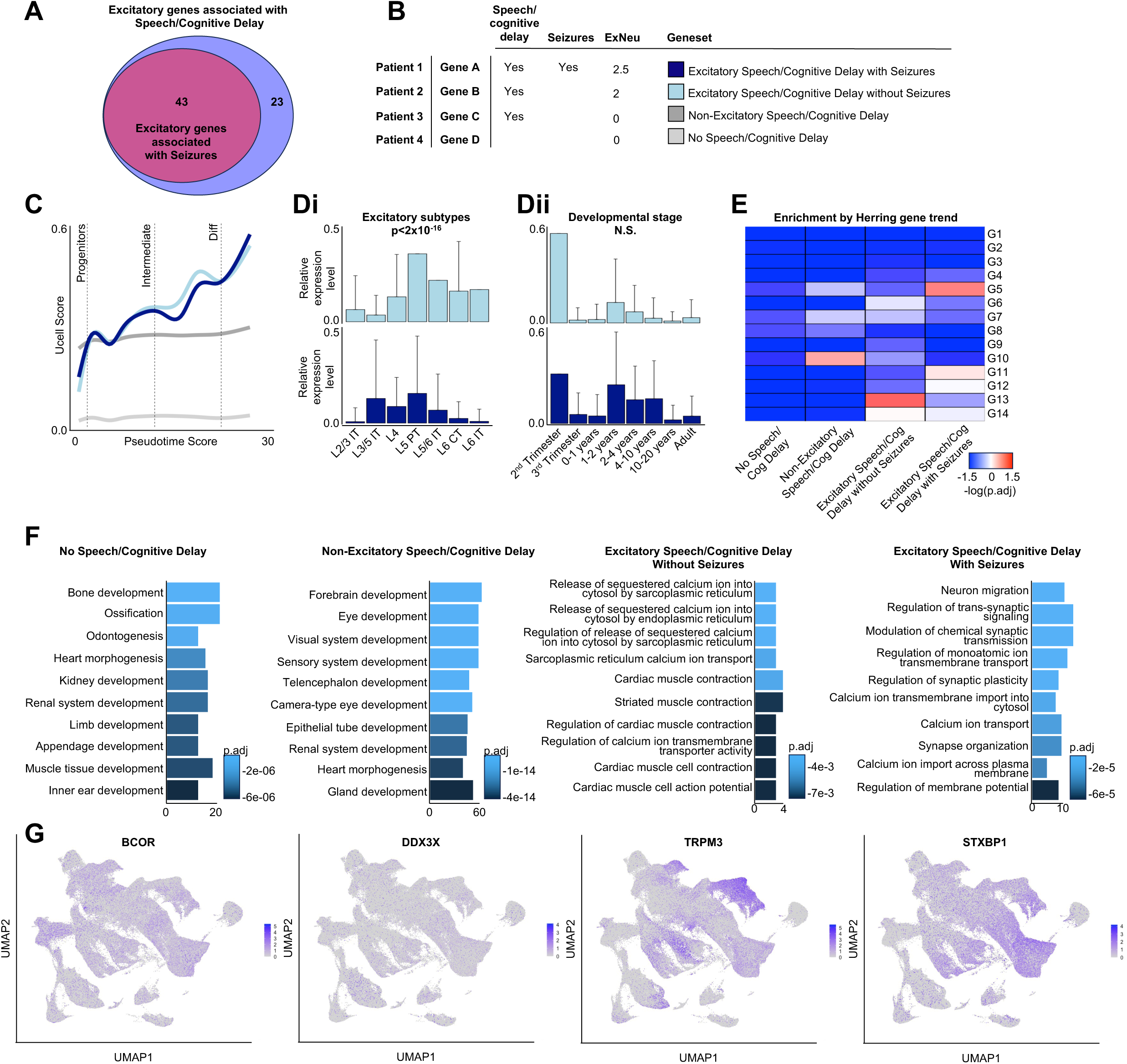
Excitatory neuron-enriched genes associated with speech/cognitive delay with or without seizures. **(A)** Excitatory neuron-enriched genes associated with seizures make up a subset (43 genes, 65%) of all excitatory neuron-enriched genes associated with speech/cognitive delay. 23 (35%) excitatory neuron-enriched genes associated with speech/cognitive delay are NOT associated with seizures. **(B)** From this data, pathogenic variants were recategorized into four new genesets as illustrated in the toy example: variants that were not associated with speech/cognitive delay (“No Speech/Cognitive Delay,” light gray), variants that were associated with speech/cognitive delay but not in genes enriched in excitatory neurons (“Non-Excitatory Speech/Cognitive Delay,” dark gray), variants that in genes enriched in excitatory neurons and associated with speech/cognitive delay but NOT seizures (“Excitatory Speech/Cognitive Delay without Seizures,” light blue), and variants in genes enriched in excitatory neurons and associated with both speech/cognitive delay AND seizures (“Excitatory Speech/Cognitive Delay with Seizures,” dark blue). **(C)** Individual cells from Velmeshev et al., 2023 demonstrated increasing aggregate geneset expression (UCell score) with increasing excitatory neuron differentiation (pseudotime score) for the two genesets of excitatory-enriched genes associated with speech/cognitive delay with or without seizures. Cells demonstrated lower aggregate expression of the Non-Excitatory Speech/Cognitive Delay and No Speech/Cognitive Delay genesets with less dramatic change over developmental time. **(Di)** Excitatory-enriched genes associated with speech/cognitive delay without seizures demonstrated higher relative gene expression in deeper cortical layers, particularly extratelencephalic neurons, whereas those associated with seizures showed higher relative expression in layer 3 -5 IT neurons (ANOVA effect by geneset p<2×10^-16^; ANOVA effect by cortical layer p<2×10^-16^; L3/5 IT across genesets p.adj=0.008; L5 PT across genesets p.adj=3.5×10^-8^; L6 CT across genesets p.adj=1.9×10^-6^). **(Dii)** Excitatory-enriched genes associated with speech/cognitive delay without seizures demonstrated a similar distribution of relative gene expression across age ranges (ANOVA effect by geneset p=0.07; ANOVA effect by age range p<2×10^-16^). **(E)** Hypergeometric testing of each of the four new speech/cognitive delay genesets shows differential enrichment patterns across “gene trend” categories G1-G14 as defined in Herring et al., 2022 (Supplemental Table 2). No enrichment remained significant after correction for multiple comparisons, however Non-Excitatory Cognitive/Speech Delay showed highest overlap with category G10 (uncorrected p=0.004), Excitatory Speech/Cognitive Delay without Seizures showed highest overlap with category G13 (uncorrected p=0.001), and Excitatory Speech/Cognitive Delay with Seizures showed highest overlap with category G5 (uncorrected p=0.003). **(F)** Gene Ontology analysis of each of the four speech/cognitive delay genesets demonstrated differential molecular pathway enrichment. In particular, Excitatory Speech/Cognitive Delay without Seizures was enriched for terms related to calcium ion transport and muscle contraction while Excitatory Speech/Cognitive Delay with Seizures was enriched for terms related to neuronal migration and trans-synaptic signaling. **(G)** Feature plots showing the expression of representative genes from each of the four genesets mapped to the UMAP of data from Velmeshev et al., 2023 (as shown in Figure 1B). Abbreviations: “Diff” = differentiated neurons; N.S. = not significant; IT = intratelencephalic neurons; PT = pyramidal tract neurons; CT = corticothalamic neurons; UMAP = uniform manifold approximation and projection; p.adj = adjusted p-value.

Based on this finding, we created four new genesets (Figure 3B; Supplemental Table 5): 1) “No Speech/Cognitive Delay” - variants in genes not associated with speech/cognitive delay, 2) “Non-Excitatory Speech/Cognitive Delay” - variants in genes associated with speech/cognitive delay that had low or no expression in excitatory neurons, 3) “Excitatory Speech/Cognitive Delay without Seizures” - variants in genes associated with speech/cognitive delay *without* seizures that were expressed in excitatory neurons, and 4) “Excitatory Speech/Cognitive Delay with Seizures” - variants in genes associated with speech/cognitive delay *and* seizures that were expressed in excitatory neurons.

To refine our expression analysis across these four categories, we calculated a UCell score for each of the four speech/cognitive delay genesets for each cell. UCell is a recently published method for evaluating aggregated geneset enrichment in individual cells of a snRNAseq dataset.^29^ As expected, UCell scores were generally higher in excitatory neurons for the two excitatory-enriched genesets (Figure 3C). Unlike the genesets not associated with speech/cognitive delay or enriched in excitatory neurons, UCell scores for the excitatory-enriched genesets increased with increasing pseudotime, implying a higher aggregate expression of genes associated with speech/cognitive delay and/or seizures with increasing excitatory neuron differentiation (Figure 3C).

We next investigated whether excitatory neuron expression biases for each of the excitatory-enriched genesets extended to more granular excitatory neuron subtypes. Cortical excitatory neurons can be further classified as intratelencephalic (IT), projecting within the cortex and striatum, or extratelencephalic (ET), projecting beyond the telencephalon to the brainstem and spinal cord.^31, 32^ IT neurons span cortical laminae but are most abundant in the upper layers, while ET neurons, which include layer 5 pyramidal tract (PT) and layer 6 corticothalamic (CT), neurons reside in deeper cortical layers. In our analysis, genes associated with speech/cognitive delay *without* seizures showed higher expression in ET neurons and other deep layer subtypes, while the expression of genes associated with speech/cognitive delay *with* seizures showed a distribution skewed toward IT neurons in layers 3-5 (ANOVA effect by geneset p<2×10^-16^; ANOVA effect by cortical layer p<2×10^-16^; Tukey L3/5 IT across genesets p.adj=0.008; Tukey L5 PT across genesets p.adj=3.5×10^-8^; Tukey L6 CT across genesets p.adj=1.9×10^-6^; Figure 3Di).

We also investigated whether excitatory-enriched genesets could be distinguished by their expression across developmental age ranges. In this case, genes associated with speech/cognitive delay with or without seizures showed bimodal expression patterns with the highest relative expression in the 2^nd^ trimester of gestation and a second protracted period of increased expression throughout childhood (Figure 3Dii). Comparing the age range distributions of these two datasets did not reveal an interaction between geneset and age range (ANOVA by geneset pval=0.07).

Herring et al., 2022 defined 14 categories of “gene trends,” which group differentially expressed genes across developmental stages by similarities in their temporal expression profiles (Supplemental Table 2).^25^ For each of the four genesets related to speech/cognitive delay outlined above, we performed hypergeometric testing to determine whether these genesets were enriched for specific Herring-defined gene trends. Although they did not remain significant after Bonferroni correction for multiple comparisons, each of the four speech or cognitive delay genesets showed distinct patterns across the different gene trend categories (Figure 3E). “No Speech/Cognitive Delay” genes were not enriched for any gene trend. “Non-Excitatory Speech/Cognitive Delay” genes were most enriched in gene trend G10, defined by high fetal expression with rapid downregulation by infancy, which includes representative genes related to chromatin organization, histone methylation, and mRNA processing. “Excitatory Speech/Cognitive Delay without Seizures” genes were most enriched in gene trend G13, defined by high fetal expression and slow downregulation by childhood, which includes representative genes related to cell death, neuron projection guidance, and locomotion. Finally, “Excitatory Speech/Cognitive Delay with Seizures” genes were most enriched in gene trend G5, defined by expression that increases from the fetal period to childhood, peaks in childhood, and then remains at a sustained level through adulthood, and contains representative genes related to axon development, trans-synaptic signaling, and learning (Figure 3E).

Unpacking these molecular categories further, gene ontology (GO) analysis suggested that genes not associated with speech or cognitive delay tended to involve pathways involved in non-CNS embryogenesis (Figure 3F,G). Genes associated with speech/cognitive delay that have low or no expression in excitatory neurons involve molecular pathways related to early processes of brain and eye development (Figure 3F,G). Genes associated with speech/cognitive delay *without* seizures involved pathways related to calcium regulation, particularly within muscle (Figure 3F,G). Finally, genes associated with speech/cognitive delay *and* seizures were more likely to involve pathways related to neuronal migration, synaptogenesis, and synaptic signaling (Figure 3F,G).

In addition to high excitatory neuron expression biases, analyses from both the Velmeshev, 2023 and Herring, 2022 datasets showed that genes associated with speech/cognitive delay also demonstrate expression biases in microglia. We investigated the relationship of microglia-biased genes with excitatory-biased genes and found that they were largely non-overlapping (Figure 4A). Of 112 unique genes associated with a phenotype of speech/cognitive delay, 63 genes (56%) exhibit an excitatory neuron expression bias, 46 genes (41%) exhibit a microglia expression bias, and 3 genes (approximately 3%) are biased toward relatively higher expression in both cell types (Figure 4A). Gene ontology analysis of microglia-biased genes associated with speech/cognitive delay included mitosis, apoptosis, TGFβ signaling, and skeletal morphogenesis (Figure 4B). By examining expression patterns of individual microglia-biased genes, we found that they largely fell into one of two categories: 1) “Micro-broad” - broad expression across greater than five percent of other cell types but with higher relative microglial enrichment, and 2) “Micro-narrow” - expression largely restricted to microglia with less than five percent expression in other cell types. Performing hypergeometric testing for enrichment of these subcategories in the Herring-defined gene trends, we found that the “Micro-broad” genes show highest enrichment for G10, a category defined by high fetal expression and rapid downregulation with representative genes involved in chromatin regulation (Figure 4C,D). Although it did not remain significant after Bonferroni correction for multiple comparisons, “Micro-narrow” genes showed highest enrichment for gene trend category G1, defined by an expression peak during infancy, with representative genes involved in receptor clustering, synapse organization, and synaptic signaling (Figure 4C,D).

**Figure 4:**
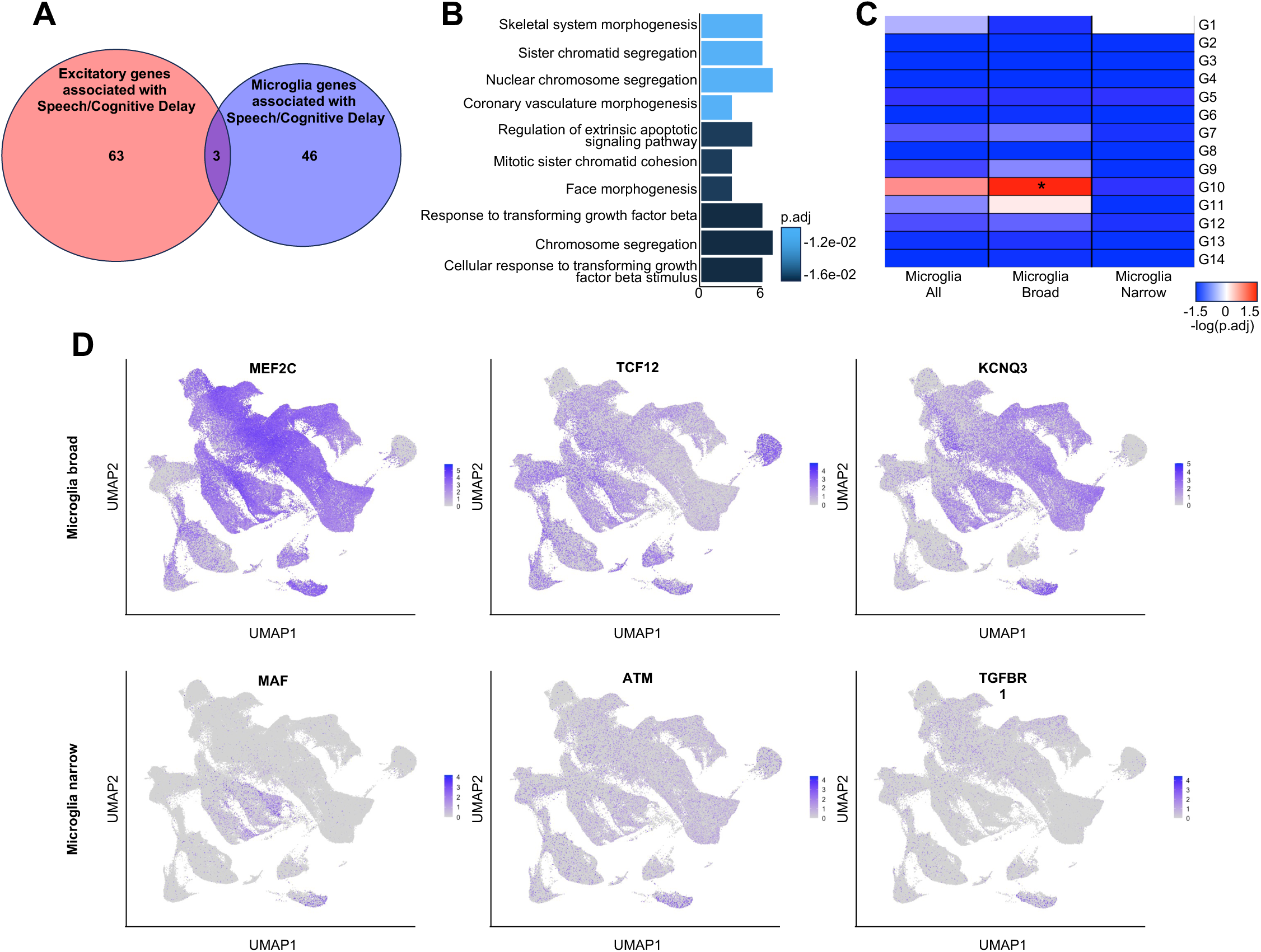
Microglia-enriched genes associated with speech/cognitive delay. **(A)** Pathogenic variants associated with speech/cognitive delay in genes with a microglial expression bias are largely non-overlapping with those that have an excitatory neuron expression bias (46 genes, 94% of microglia-enriched genes associated with speech/cognitive delay). **(B)** Gene Ontology analysis of the 49 microglia-enriched genes associated with speech/cognitive delay reveals molecular pathways related to chromosome segregation during mitosis and transforming growth factor beta (TGFβ) signaling. **(C,D)** Microglia-enriched genes associated with speech/cognitive delay could be subdivided into two genesets, “Microglia Broad” and “Microglia Narrow,” depending on whether genes were expressed in greater or less than 5% of non-microglia. Hypergeometric testing of microglial genesets with “gene trends” G1-G14, as defined in Herring et al., 2022 (Supplemental Table 2), demonstrated that Microglia Broad genes were significantly enriched for gene trend G10 (adjusted pval=0.03), while Microglia Narrow genes did not show significant enrichment after correction for multiple comparisons but were most enriched for gene trend G1 (uncorrected pval=0.02, corrected pval=0.97; **C**). Feature plots **(D)** showing the expression of representative genes from Microglia Broad and Microglia Narrow genesets mapped to the UMAP of data from Velmeshev et al., 2023 (as shown in Figure 1B). Abbreviations: UMAP = uniform manifold approximation and projection; p.adj = adjusted p-value. * = p<0.05.

Finally, we investigated whether certain phenotype-related genesets might show a developmental stage-specific expression bias, rather than a cell type-specific expression bias. The relative expression and breadth of expression for genes associated with seizures were biased toward infancy and childhood in both the Velmeshev, 2023 and Herring, 2022 datasets (Supplemental Figure 3), however the Velmeshev, 2023 dataset showed a wider developmental window of expression bias from the 3^rd^ trimester of gestation into adulthood. The two datasets differed in the developmental expression biases of genes associated with speech/cognitive delay: the Herring, 2022 dataset suggested expression biases toward the fetal/neonatal period, whereas the Velmeshev, 2023 dataset suggested expression biases from two to four years old. Additionally, in the Herring, 2022 dataset, there was a bias for genes associated with motor delay for the fetal and infant stages, which was not observed in the Velmeshev, 2023 dataset. Finally, although none of the variants associated with structural brain abnormalities showed cell type expression biases, variants associated with CCD did show expression biases for genes enriched in the 2^nd^ trimester in the Velmeshev, 2023 dataset (this effect was also observed in the Herring, 2022 dataset but did not remain significant after Bonferroni correction for multiple comparisons, perhaps due to the bundling of early trimesters into a single “fetal” developmental stage category; Supplemental Figure 3). This suggests that variants associated with CCD may broadly disrupt early cortical development across multiple cell types. Indeed, gene ontology analysis for genes associated with CCD that also show 2^nd^ trimester expression biases resulted in pathways associated with both neuroepithelial and glial differentiation, as well as maintenance of the blood brain barrier and synapse organization (data not shown).

## Discussion

Macroanatomic structural brain malformations and microstructural functional NDDs together contribute to a high burden of pediatric disability;^2,3,6, 9^ however, the increasing availability and sophistication of clinical genetic testing has led to an improved detection of genetic etiologies for many patients with NDDs.^15, 16, 19, 20^ While this has yielded a more detailed understanding of genotype-phenotype relationships for many NDDs, our ability to model, perturb, and ultimately disentangle the cellular pathophysiology of NDDs requires us to first understand the cellular substrates that mediate neurodevelopmental symptoms. This requires a link from genotype to cell type to phenotype. We set out to investigate this link by capitalizing on large scale neurodevelopmental phenotype data and merging it with the most comprehensive snRNAseq datasets from developing human cortex to date in order to identify cell-specific expression biases of genes associated with specific neurodevelopmental phenotypes. The goals of our current study were four-fold: 1) To combine large scale pathogenic variants from multiple neurodevelopmental phenotypes with snRNAseq data, allowing us to incorporate a higher frequency of rare variants, to reduce the bias associated with curated lists of variants, and to improve our power to detect shared patterns of gene expression within broad phenotypes; 2) To enhance our confidence in cell-specific gene expression signatures by using massive, recently generated snRNAseq datasets of approximately 150,000 to 350,000 cells with higher depth of sequencing and more comprehensive inclusion of rare cortical cell types; 3) To begin to build hypotheses about the cortical pathophysiology that mediates the cellular link between genotype and neurodevelopmental phenotype; and finally, 4) To demonstrate how large scale, cross-modality datasets from existing repositories can be integrated and reanalyzed to draw novel conclusions about the pathophysiology of NDDs.

We found that pathogenic single gene variants associated with speech/cognitive delay, as well as pathogenic single gene variants associated with seizures, showed enhanced expression by multiple metrics in excitatory cortical neurons, a finding that was reproduced in two separate human snRNAseq datasets. Single gene variants associated with speech/cognitive delay also showed enhanced expression in microglia, an expression bias that was not observed for genes associated with seizures. Conversely, subjects whose pathogenic variant affected a gene enriched in excitatory neurons or microglia were significantly more likely to have speech/cognitive delay, while seizures were more likely only for subjects with a pathogenic variant in a gene enriched in excitatory neurons.

Excitatory-enriched genes associated with seizures comprised a subset of those associated with speech/cognitive delay, suggesting that variants in excitatory-enriched genes from subjects with speech/cognitive delay can be parsed into two phenotypic subsets. Both excitatory-enriched genesets showed continual increases in expression throughout neuronal differentiation, as illustrated by increasing UCell scores, a measure of aggregate geneset expression per cell, with pseudotime, a proxy for differentiation previously calculated in Velmeshev et al., 2023.^26^ In contrast, aggregate geneset expression did not increase dramatically with excitatory neuron differentiation for genes that were not associated with speech/intellectual disability or for genes that did not demonstrate expression biases in excitatory neurons. Despite sharing high expression in differentiated excitatory neurons, there were some key differences in molecular characterization among excitatory-enriched genes associated with speech/cognitive delay with or without seizures.

Excitatory-enriched genes associated with speech/cognitive delay *without* seizures tended to involve molecular pathways related to calcium regulation and showed their highest expression bias in deep cortical layers - primarily extratelencephalic neurons. These genes also showed maximal enrichment in a Herring et al., 2022 gene trend class that encompasses genes related to neurite development, cell death, and locomotion. This Herring et al., 2022 gene trend class also exhibits peak expression in fetal brain development followed by slow downregulation, similar to the high second trimester expression bias we observed when comparing genesets across developmental stage. In this subset, pathogenic variants affected genes encoding transcription factors such as *SOX5*, *PBX1*, and *MEIS2*, which have been identified in individuals with intellectual disability and developmental delay, typically without seizures. It is also notable that the Herring et al., 2022 gene trend category includes “locomotion,” and that our GO terminology includes terms related to muscle contraction and calcium transport in the sarcoplasmic reticulum, since representative genes from this geneset also include *DMD* and *RYR2*. Although many of these disorders are classically considered neuromuscular, there is an increasing appreciation for the non-muscular central nervous system impact of these disorders that results in developmental delay for some patients.^33, 34^

In contrast, pathogenic variants in excitatory-enriched genes associated with both speech/cognitive delay *and* seizures involved molecular pathways related to neuron migration and synaptic communication. These genes showed more even expression among intratelencephalic and extratelencephalic subsets of excitatory neurons, with highest expression in layer 3/5 IT neurons, as well as layer 5 PT neurons. These genes also demonstrated highest overlap with a Herring et al., 2022 gene trend category of genes related to synaptic signaling. Although this geneset showed the highest expression bias in the second trimester of gestation, perhaps reflecting early expression of transcription factors (*DCX*, *CUX2*, *FOXG1)* and cytoskeletal genes (*TUBB*, *TUBA1A*, *TUBB2B*, *ACTB*, *PHACTR1*), genes from the Herring et al., 2022 gene trend category are described to increase in expression from the fetal period through childhood and remain sustained through adulthood, likely reflective of synaptic regulatory genes and ion channel subunits crucial for lifelong trans-synaptic signaling (*CNKSR2*, *KCNQ3*, *GABRB3*). This pattern is reflected in the bimodal distribution of developmental stage-specific enrichment that increases again in childhood stages.

Finally, although driven more by rare variants than for excitatory neurons, pathogenic variants associated with speech/cognitive delay also involved a set of microglia-enriched genes that were largely distinct from those enriched in excitatory neurons. This is of particular note, given our expanding appreciation for the role microglia play in synaptic development and nervous system pathophysiology.^35, 36^ Microglia-enriched genes were not necessarily expressed *only* within microglia, but rather showed enhanced relative expression in microglia, involving pathways related to chromosome regulation during mitosis. For example, *ARID1A* and *ARID1B*, which are associated with a phenotype of Coffin-Siris syndrome that exhibits more frequent learning and cognitive delay,^37–39^ or *ATM*, which causes ataxia-telangiectasia in part through ATM-dependent microglial dysregulation,^40^ all show higher relative expression in microglia than other cortical cell types. It is therefore compelling to hypothesize that neurodevelopmental dysfunction is a result of microglial cell cycle dysregulation that in turn results in aberrant neuronal circuit maturation. Perhaps a more realistic hypothesis for genes enriched in but not specific to microglia is that neuropathology is the result of both neuronal dysfunction and aberrant microglial synaptic refinement.^41^ At a minimum, this analysis uncovers enriched gene expression in additional cortical cell types that may have been previously underappreciated in our understanding of the pathophysiology of Coffin-Siris syndrome, ataxia telangiectasia, and other NDDs.

As intriguing as the neurodevelopmental phenotypes that did show cell-specific expression biases are those that did *not* show cell-specific or stage-specific expression biases. For example, across all analyses we did not find a cell type-specific expression bias for autism-associated genes, which may argue against convergence of autism associated genes in one or a few cortical cell types.^42^ However, despite broad cell type expression, a recent study using long-read sequencing in human cortex suggests that autism-associated genes tend to have a wide variety of splice isoforms that may each have differential impacts on cortical cell types.^24^

Likewise, we did not find cell type enrichment for genes associated with hydrocephalus/ventriculomegaly, unlike Duy et al., 2022 who mapped genes associated with congenital hydrocephalus into human cortical single cell transcriptomic data and reported enrichment in excitatory cortical neurons and their neuroepithelial precursors.^28^ Discrepancies in cell type-specific expression may result from inclusion criteria of the initial cohort; Duy et al., 2022 limited their analysis to subjects with congenital hydrocephalus whose symptoms were severe enough to warrant neurosurgical cerebrospinal fluid (CSF) diversion,^28^ whereas our cohort included patients with any form of symptomatic or asymptomatic ventriculomegaly with or without associated hydrocephalus.

Strengths of our study include the massive scale of phenotypic and human cortical snRNAseq datasets utilized in our analysis. We were able to improve the robustness of our findings by reproducing them in two separate recently published snRNAseq from human cortex across development.^25, 26^ Coupling the large scale of the DDD genotype-phenotype dataset with our single-institution data collected from neonatal subjects with a greater proportion of congenital brain malformations powered us to analyze a broad range of NDDs across developmental stages. Moreover, unlike studies that perform enrichment analyses using curated genesets,^22^ our analysis utilizes a more unbiased approach to phenotypic geneset categorization based on any presenting phenotype, rather than solely a classical neurodevelopmental phenotype, allowing genes to be grouped in more than one phenotypic geneset, including both negative and positive phenotypic genesets if that represented the range of presentations in our cohort. Finally, we demonstrate an analytic pipeline that repurposes existing data from publicly accessible repositories. Rather than continually and laboriously generating new datasets, much can be inferred about cell type expression biases by reanalyzing the wealth of existing genotype, phenotype, and transcriptomic data generated by the modern expansion of cutting edge clinical and scientific genetic profiling techniques.

A major limitation of our study is its sole reliance on single nucleus transcriptomic data, which does not incorporate the myriad other modulatory genetic mechanisms that regulate neurodevelopment. As a result, we can only analyze pathogenic variants that putatively directly alter exons of individual genes, precluding the analysis of aneuploidies, large copy number variations, epigenetic alterations, non-coding regions, and the modifying effects of combinations of variants that might contribute to an oligogenic phenotype.^43^ Additionally, built into the use of transcriptomic data is the assumption that the level of gene expression in each cell type correlates with that cell’s degree of dysfunction from a pathogenic variant, an assumption that is highly unlikely to be true across all genes included in this study. However, at least at a fundamental level, a cell that does not express a pathogenic variant at all is probably less likely to be affected than a cell that does express the variant, which is why we attempted to coarsely correct for this possibility by binning genes into those with positive expression versus no/low expression within each cell type. Finally, our study focused on human cortical snRNAseq datasets, given the predominant role the cortex is likely to play in mediating NDDs.^12, 27, 28, 44^ However, repeating this analysis in striatal, cerebellar, or brainstem datasets would almost certainly uncover broader cell type specific expression biases across the nervous system.

The goal of this study was to begin linking phenotype-associated genes with cortical cell type pathophysiology, and while we find that genes associated with speech/cognitive delay and/or seizures are enriched in excitatory cortical neurons and microglia, additional research will be necessary to validate convergent and divergent molecular pathways, elucidate how cell type-specific cortical pathophysiology functionally mediates neurodevelopmental symptoms, and elaborate on our findings by incorporating other modalities of genetic regulation. For example, other existing multi-modal “omics” repositories could be harnessed to investigate cell-specific chromatin accessibility of gene regulatory regions^45, 46^ or splice isoforms,^24^ or to correlate cell specific-protein translation with our predictions around cell-specific transcription^47^ for genes associated with specific neurodevelopmental phenotypes. Finally, although large scale transcriptomic analyses like this one highlight new hypotheses and areas of focus, they do not obviate the need for detailed functional molecular studies of individual genes that determine how pathogenic variants affect protein function, and how protein dysfunction in turn leads to cellular pathology, alters developmental mechanics, disrupts neuronal microcircuitry, and ultimately generates neurodevelopmental symptomatology. To this end, the union between the increased availability of high-depth clinical genetic testing,^48^ the increasing scale of multimodal single cell profiling,^31, 49^ and novel methods for testing the impact of pathogenic variants on human cortical development^43^ via organotypic slice culture,^50^ cerebral organoids,^51–53^ and humanized animal models,^54^ will together help unlock the complexity of human NDDs.

## Acknowledgements

The authors would like to thank Dr. Dmitry Velmeshev for his thoughtful comments on this manuscript.

---

**Supplemental Figure 1:**
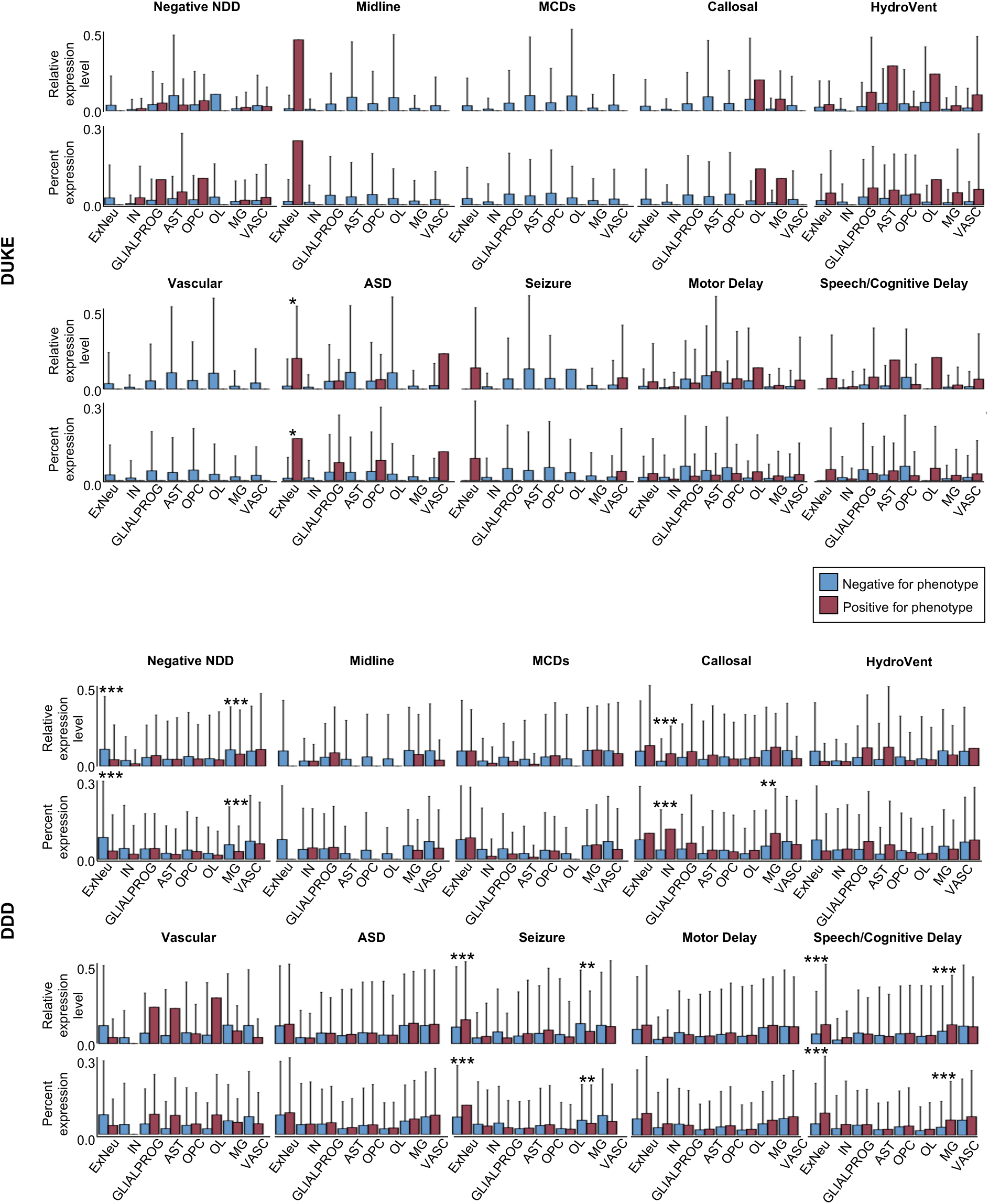
Cell-specific expression biases separated by Duke versus DDD dataset. Reanalysis of the cell-specific expression biases in the Velmeshev et al., 2023 human cortical snRNAseq data shows that significant expression biases are driven mainly by the more highly powered DDD data. However, the primary findings of higher expression in excitatory neurons of genes associated with seizures and speech/cognitive delay still demonstrate a similar directionality in the Duke dataset. Interestingly, the only significant expression bias isolated to the Duke dataset is excitatory neuron expression bias of autism-associated genes. Red columns = genes associated with the phenotype; blue columns = genes not associated with the phenotype. Abbreviations: NDD = neurodevelopmental disorder; Midline = midline defects; MCDs = malformations of cortical development; Callosal = callosal abnormalities; HydroVent = hydrocephalus/ventriculomegaly; Vascular = vascular disorders, hemorrhage, or stroke; ASD = autism spectrum disorder; ExNeu = excitatory neurons; IN = inhibitory neurons; GLIALPROG = glial progenitors; AST = astrocytes; OPC = oligodendrocyte precursor cells; OL = oligodendrocytes; MG = microglia; VASC = endothelial cells/pericytes. * = p<0.05; ** = p<0.01; *** = p<0.001.

**Supplemental Figure 2:**
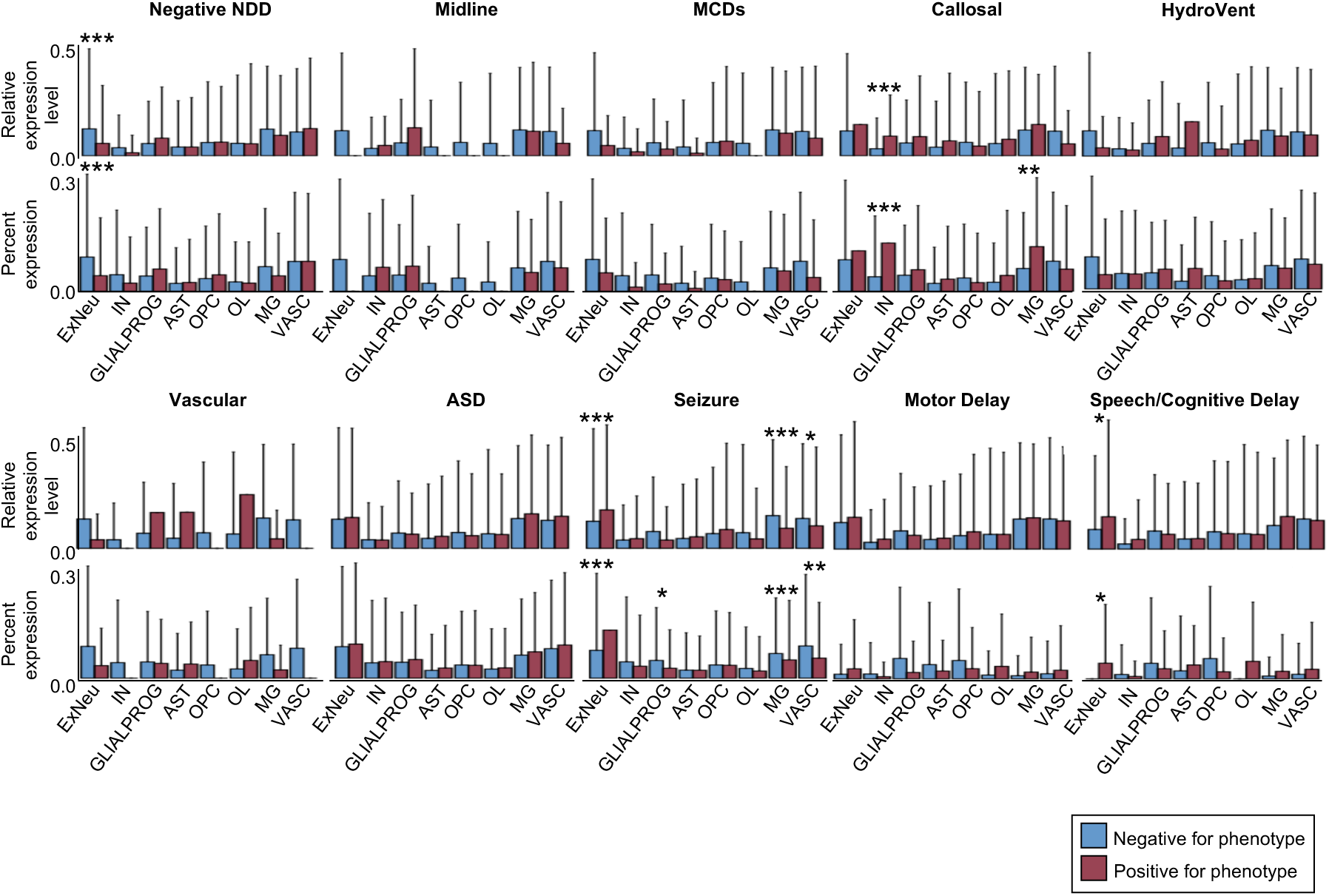
Cell-specific expression biases excluding genes associated with rare pathogenic variants. Mean relative expression level (top; log2 fold change in expression compared to all other cell types) and mean percent expression (bottom; percent of cell type expressing genes of interest) analyzed in nuclei from Velmeshev et al., 2023 with phenotype-related genesets that exclude genes in which pathogenic variants were observed in fewer than five subjects. Compared to the complete analysis in Figure 1, expression bias in microglia of genes associated with speech/cognitive delay is no longer observed, suggesting that this effect is driven primarily by genes associated with rare variants. Red columns = genes associated with the phenotype; blue columns = genes not associated with the phenotype. Abbreviations: NDD = neurodevelopmental disorder; Midline = midline defects; MCDs = malformations of cortical development; Callosal = callosal abnormalities; HydroVent = hydrocephalus/ventriculomegaly; Vascular = vascular disorders, hemorrhage, or stroke; ASD = autism spectrum disorder; ExNeu = excitatory neurons; IN = inhibitory neurons; GLIALPROG = glial progenitors; AST = astrocytes; OPC = oligodendrocyte precursor cells; OL = oligodendrocytes; MG = microglia; VASC = endothelial cells/pericytes. * = p<0.05; ** = p<0.01; *** = p<0.001.

**Supplemental Figure 3:**
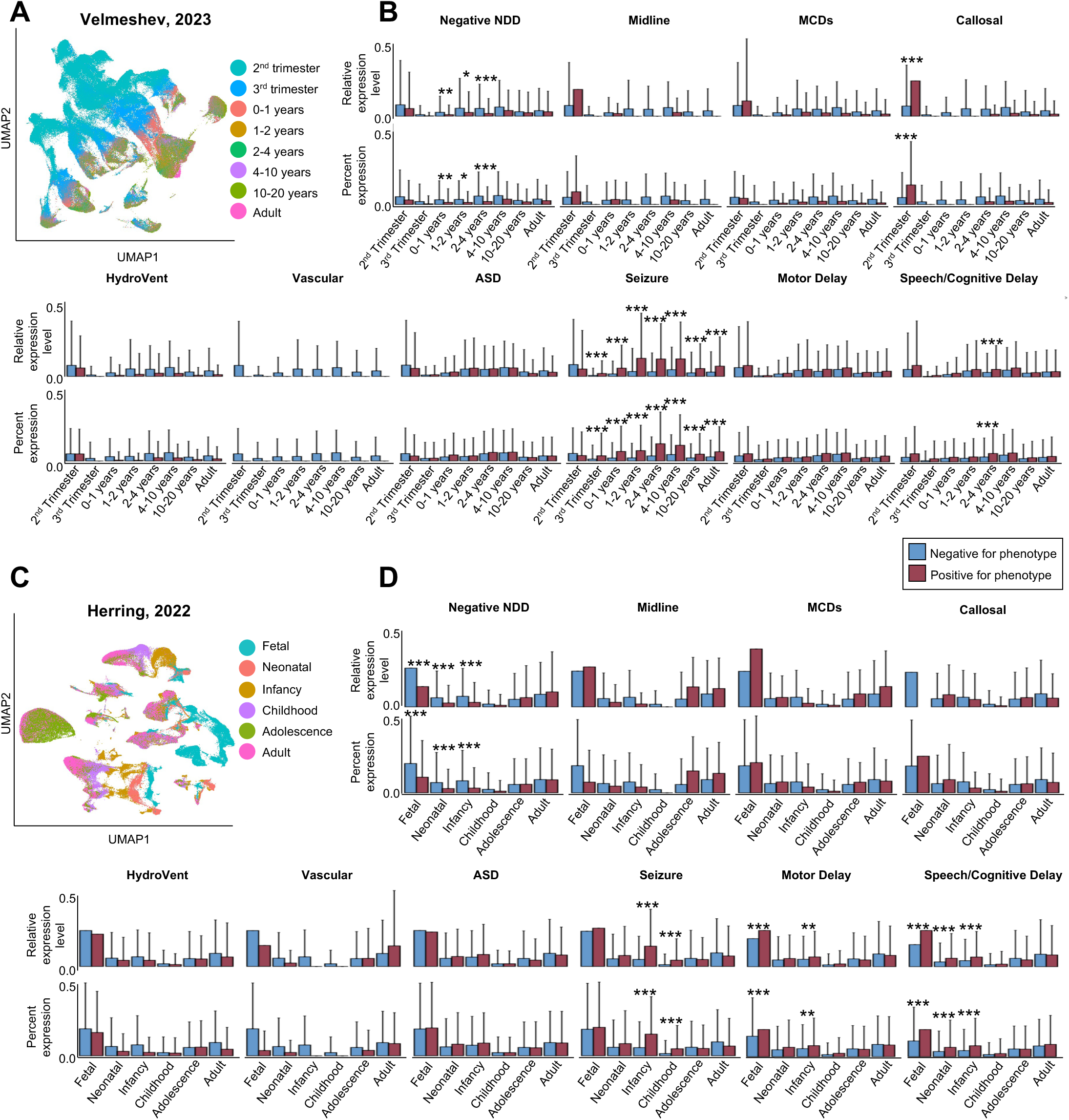
**(A)** UMAP from Velmeshev et al., 2023 with originally assigned age range annotations. **(B)** Mean relative expression level (top; log2 fold change in expression compared to all other age ranges) and mean percent expression (bottom; percent of cells in each age range expressing genes of interest) demonstrate age range-specific expression biases in human cortex. **(C)** UMAP from Herring, et al., 2022 with originally assigned age range annotations. **(D)** Mean relative expression level (top) and mean percent expression (bottom) demonstrate somewhat discrepant age range-specific expression biases from those observed in **(B)**. In **(B)**, seizure-related genes show a wider expression bias across all age ranges after 2^nd^ trimester, whereas in **(D)** seizure-related genes show a more restricted expression bias in infancy and childhood. Moreover, in **(B)**, genes associated with speech/cognitive delay only show a significant expression bias around 2 to 4 years old, but in **(D)** they show a bias toward nuclei from the fetal stage through infancy. Red columns = genes associated with the phenotype; blue columns = genes not associated with the phenotype. Abbreviations: NDD = neurodevelopmental disorder; Midline = midline defects; MCDs = malformations of cortical development; Callosal = callosal abnormalities; HydroVent = hydrocephalus/ventriculomegaly; Vascular = vascular disorders, hemorrhage, or stroke; ASD = autism spectrum disorder; ExNeu = excitatory neurons; IN = inhibitory neurons; GLIALPROG = glial progenitors; AST = astrocytes; OPC = oligodendrocyte precursor cells; OL = oligodendrocytes; MG = microglia; VASC = endothelial cells/pericytes; UMAP = uniform manifold approximation and projection. * = p<0.05; ** = p<0.01; *** = p<0.001.

**Supplemental Table 1: Human phenotype ontology (HPO) terms used to annotate DDD phenotypes**

**Supplemental Table 2: Gene trend annotations reproduced from Herring et al., 2022**

**Supplemental Table 3: Phenotype specific genesets for Duke and DDD datasets organized by geneset**

**Supplemental Table 4: Phenotype specific genesets for Duke and DDD datasets organized by gene**

**Supplemental Table 5: Speech/cognitive delay genesets and microglia-enriched genesets**

